# Identification of feeding apparatus components in a heterotrophic marine flagellate

**DOI:** 10.64898/2026.03.30.714256

**Authors:** Gillian Clifford, Samuel J. P. Taylor, Midori Ishii, Fernanda Cisneros-Soberanis, Bungo Akiyoshi

## Abstract

Acquiring nutrients is a fundamental biological process of all organisms, playing crucial roles in ecological sustainability. Diplonemids are highly abundant heterotrophic unicellular flagellates that are widespread in the world’s ocean. They have a highly complex microtubule-based feeding apparatus (cytostome-cytopharynx complex) located adjacent to the deep flagellar pocket from which two flagella emerge from parallel basal bodies. The apical papilla is a tongue-shaped structure unique to diplonemids that connects the cytopharynx and the flagellar pocket, the latter of which is formed by reinforcing microtubules (MTR) and two flagellar roots called intermediate and dorsal roots. Here we report identification of 17 proteins that localize at the feeding apparatus or flagellar apparatus in *Diplonema papillatum*. Using ultrastructure expansion microscopy, we show that Mad2 and its interaction partner MBP65 localize at the MTR, intermediate root, and dorsal root. Homologs of proteins that associate with the flagellar apparatus in *Trypanosoma brucei* (PFR2, KMP11, BILBO1) localize at the feeding apparatus in *D. papillatum*. We also identify proteins that localize at the apical papilla, MTR, parallel microtubule loop, or cytopharynx. By discovering components of the feeding apparatus for the first time in diplonemids, this work forms the foundation to understand molecular mechanisms of the feeding apparatus in these highly abundant marine plankton.

## Introduction

Obtaining nutrients is a fundamental, universal, and continuous activity for all living organisms with an ecological importance of the planet (Cavalier-Smith, 2013; Leander, 2020). Diplonemids are a group of heterotrophic flagellated eukaryotes that are divergent from traditional model eukaryotes (Burki et al., 2020). They are highly abundant, diverse, and widespread in the world’s ocean with an inescapable ecological significance (de Vargas et al., 2015; Gawryluk et al., 2016; Flegontova et al., 2016, 2020; Schoenle et al., 2021; Tashyreva et al., 2022; Pilátová et al., 2023). Diplonemids have a highly complex microtubule-based feeding apparatus, characterized by a cytostome (cell mouth) located at the anterior end of the cell, leading into a tubular cytopharynx (gullet) (Montegut-Felkner and Triemer, 1996; Tashyreva et al., 2018; Prokopchuk et al., 2019; Tashyreva et al., 2023). The cytostome locates on the right side of the deep flagellar pocket from which two flagella emerge from basal bodies lying parallel (Porter, 1973). The apical papilla, a tongue-shaped structure unique to diplonemids, connects the cytopharynx and the flagellar pocket (Figure 1). It continues as a structure called reinforcing microtubules (MTR), which contributes to the formation of the flagellar pocket. The MTR originates underneath the flagellar pocket and also creates the flagellar pocket extension (Montegut-Felkner and Triemer, 1994).

**Figure 1.**
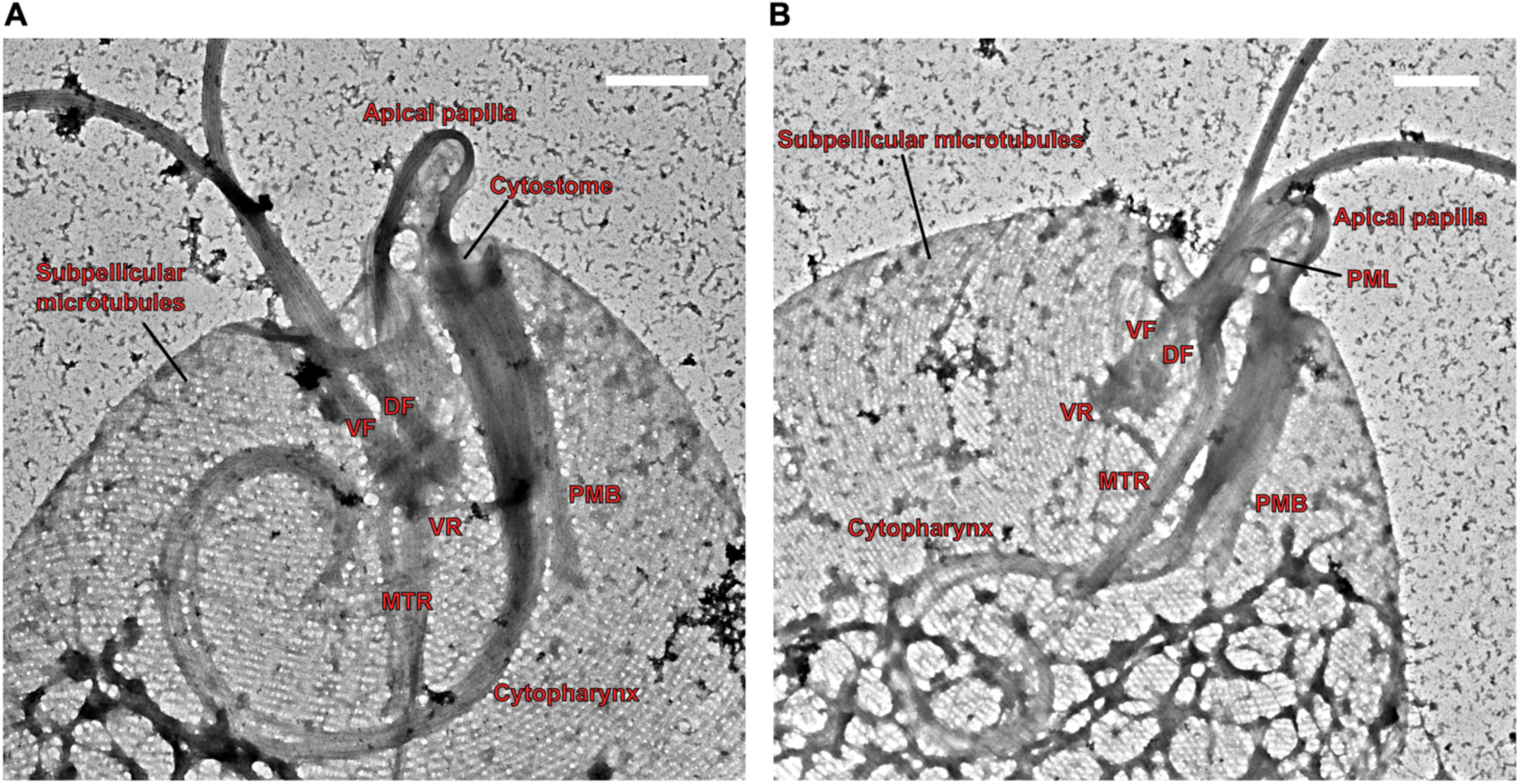
TEM analysis of detergent-extracted *Diplonema papillatum.* (A and B) The feeding apparatus locates adjacent to the flagellar apparatus. Cells settled onto a grid were treated with detergent and fixed with glutaraldehyde, followed by a positive staining with uranyl acetate. A connection is observed between the tip of MTR and cytopharynx. Note that dorsal root and intermediate root are not readily discernible due to overlap with other structures such as basal bodies and sub-pellicular microtubules. Magnification 5kX and 4kX. VF: ventral flagellum, DF: dorsal flagellum, VR: ventral root, MTR: reinforcing microtubules, PMB: peripheral microtubule band. Scale bars, 1 µm.

In many protists, flagellar roots that associate with basal bodies play various important roles such as regulation of cell shape, motility, and feeding (Moestrup, 2000). Diplonemids have three asymmetrically arranged flagellar roots: a dorsal root (DR), an intermediate root (IR), and a ventral root (VR) (Montegut-Felkner and Triemer, 1994). The DR originates from the dorsal basal body, while the IR originates from the ventral basal body and the VR is laterally associated with the ventral basal body. The flagellar pocket is reinforced by two microtubule roots (DR and IR) and MTR (which is not a flagellar root in diplonemids) (Montegut-Felkner and Triemer, 1994; Simpson, 1997). Little is known about the molecular mechanism of the formation or regulation of flagellar roots.

Diplonemids are evolutionarily closely related to kinetoplastids, which include important pathogens such as *Trypanosoma brucei* which causes African trypanosomiasis and *Trypanosoma cruzi* which causes Chagas disease (Cavalier-Smith, 2016). A feeding apparatus is present in free-living kinetoplastids (Brugerolle, 1979; Tikhonenkov et al., 2021) and some parasitic trypanosomatids such as *T. cruzi* (Alcantara et al., 2014; Chasen et al., 2019, 2020), while it is lost in many parasitic trypanosomatids including *T. brucei* and *Leishmania* (Halliday et al., 2021; Cunha-E-Silva et al., 2025). Despite the widespread presence in flagellates and ciliates, relatively little is known about the molecular components or mechanistic understanding of the cytostome-cytopharynx complex (Guerrier et al., 2017; Etheridge, 2022). In diplonemids, basically nothing is known about the composition of the feeding apparatus.

As part of our ongoing effort to dissect mitotic mechanisms in diplonemids (Akiyoshi et al., 2025), we have examined the localization of various proteins by endogenous YFP-tagging in genetically-tractable *Diplonema papillatum* (also known as *Paradiplonema/Isonema papillatum*) (Faktorová et al., 2020, 2026; Tashyreva et al., 2022; Valach et al., 2023). Here we report the identification of 17 proteins that localize to the feeding apparatus and/or flagellar apparatus.

## Results and Discussions

### TEM analysis of detergent-treated cells

We first visualized the feeding apparatus in detergent-extracted *D. papillatum* cells by transmission electron microscopy (TEM) (Figure 1). In these images, cytostome, cytopharynx, apical papilla, VR, MTR, parallel microtubule loop (PML), two flagella and basal bodies were visible. These results confirm that the feeding apparatus of *D. papillatum* is highly similar to that of *D. ambulator* (Montegut-Felkner and Triemer, 1996).

### Mad2 and MBP65 localize at flagellar roots and MTR

In traditional model eukaryotes such as yeast and human, Mad2 localizes at kinetochores and is involved in the spindle checkpoint control mechanism (Musacchio, 2015). However, in *T. brucei*, Mad2 and its interaction partner MBP65 localize at the microtubule quartet (Akiyoshi and Gull, 2013; Akiyoshi, 2020; Billington et al., 2023), which is thought to be the remnant of the DR (Lacomble et al., 2009; Yubuki et al., 2016). In *D. papillatum*, the Mad2 homolog localizes near basal bodies and the feeding apparatus (Figure 2A) (Akiyoshi et al., 2025). To examine its precise localization, we treated cells with detergent to burst the cells before imaging (Figure 2B). When the feeding apparatus was detached from the cell body, Mad2-YFP signal was found on a cytopharynx-proximal part of the MTR (Figure 2C). We next performed ultrastructure expansion microscopy (U-ExM) (Chen et al., 2015; Gambarotto et al., 2019). Immunostaining with an anti-tubulin antibody allowed us to visualize flagellar roots (Figure 2D). This analysis revealed Mad2-YFP signal on IR and DR, but not VR. Additional signal was found on the cytopharynx-proximal part of the MTR, which seemingly corresponds to the flagellar pocket extension area rather than the flagellar pocket area (Figure 2D).

**Figure 2.**
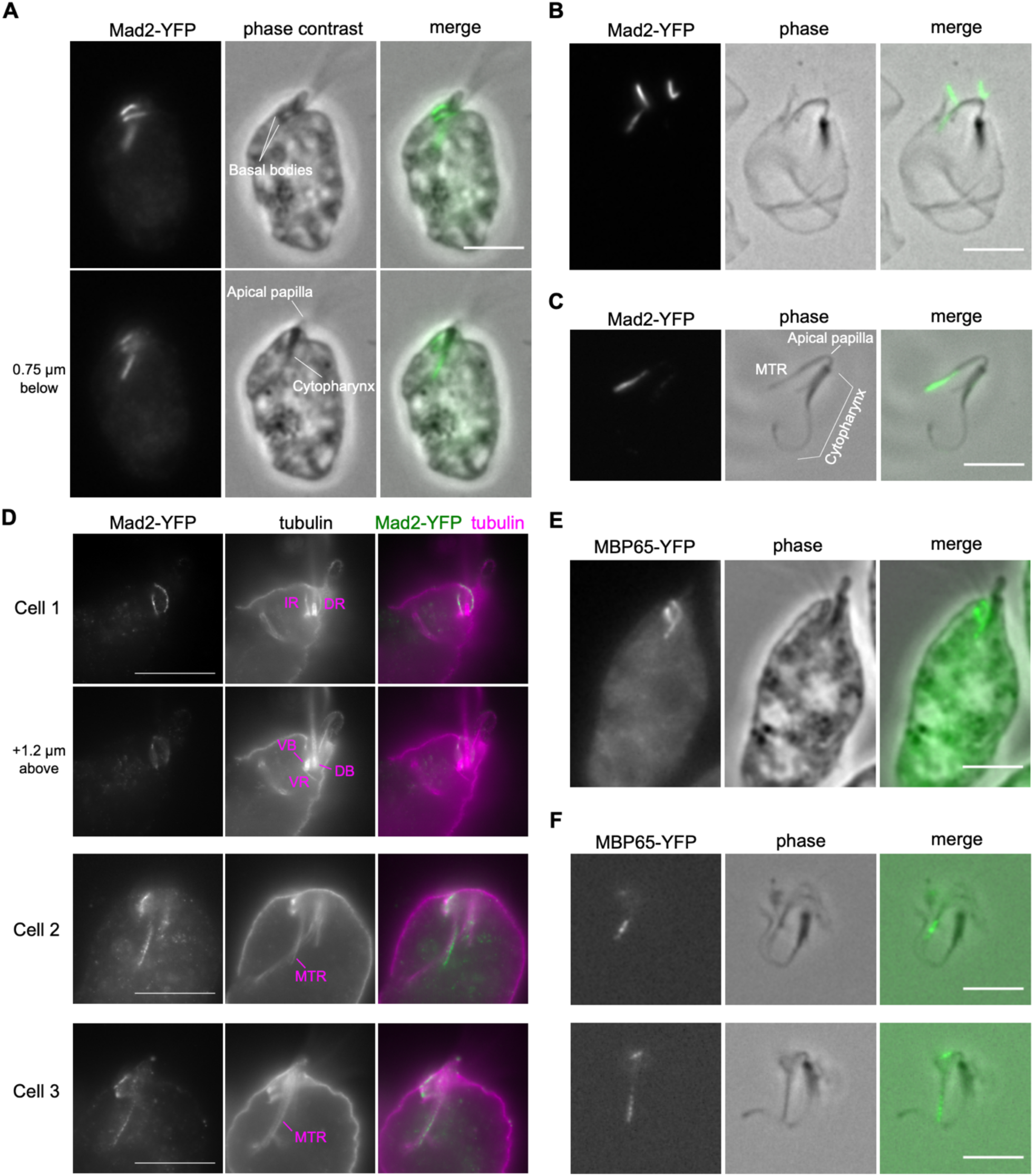
Mad2 and MBP65 localize at the intermediate root, dorsal root, and MTR. (A) Mad2-YFP localizes near basal bodies and cytopharynx, as previously reported (Akiyoshi et al., 2025). Cells were fixed with formaldehyde. Two different z sections are shown. (B, C) Mad2 localizes at part of the MTR. Cells were treated with detergent, followed by fixation with formaldehyde. (D) U-ExM of cells expressing Mad2-YFP showing its localization at the IR, DR, and MTR. Immunostaining was performed using anti-GFP antibody and anti-tubulin antibody. Two different z sections are shown for Cell 1 to highlight the three flagellar roots. The ventral basal body (VB) and dorsal basal body (DB) were determined by the presence or absence of the VR, respectively. Scale bars, 20 µm. (E, F) MBP65 has a similar localization pattern as Mad2. Cells expressing MBP65-YFP without (E) or with (F) detergent treatment were imaged. IR: intermediate root, DR: dorsal root, VR: ventral root, VB: ventral basal body, DB: dorsal basal body, MTR: reinforcing microtubules. Scale bars, 5 µm unless otherwise noted. Cell lines, DP6: Mad2-YFP, DP85: MBP65-YFP.

We next examined the localization of the MBP65 homolog and found a similar localization pattern (Figure 2E, F). These results show that Mad2 and MBP65 localize at a subset of flagellar roots in both diplonemids and kinetoplastids. In addition, they localize at the MTR in diplonemids.

### POLO1 localizes at the dorsal root, while POLO2 localizes at and near basal bodies

Polo-like kinases (PLKs) play diverse functions in eukaryotes (Zitouni et al., 2014). *D. papillatum* has two proteins that are homologous to polo-like kinases (Benz et al., 2024), named POLO1 (DIPPA_20058) and POLO2 (DIPPA_32534) herein. POLO1-YFP showed signal near one of the two basal bodies and at an unknown location (Figure 3A). U-ExM revealed that the protein is on DR, not on IR (Figure 3B). POLO2 showed complex localization patterns, including at and around basal bodies (Figure 4A, B). The POLO2 signal was not found in all cells, suggesting that it is likely cell cycle regulated. Although further work is needed to determine its precise locations or reveal functions, we speculate that POLO2 might be functionally related to PLK of *T. brucei*, which localizes at various cytoskeletal structures including basal bodies (Lozano-Núñez et al., 2013).

**Figure 3.**
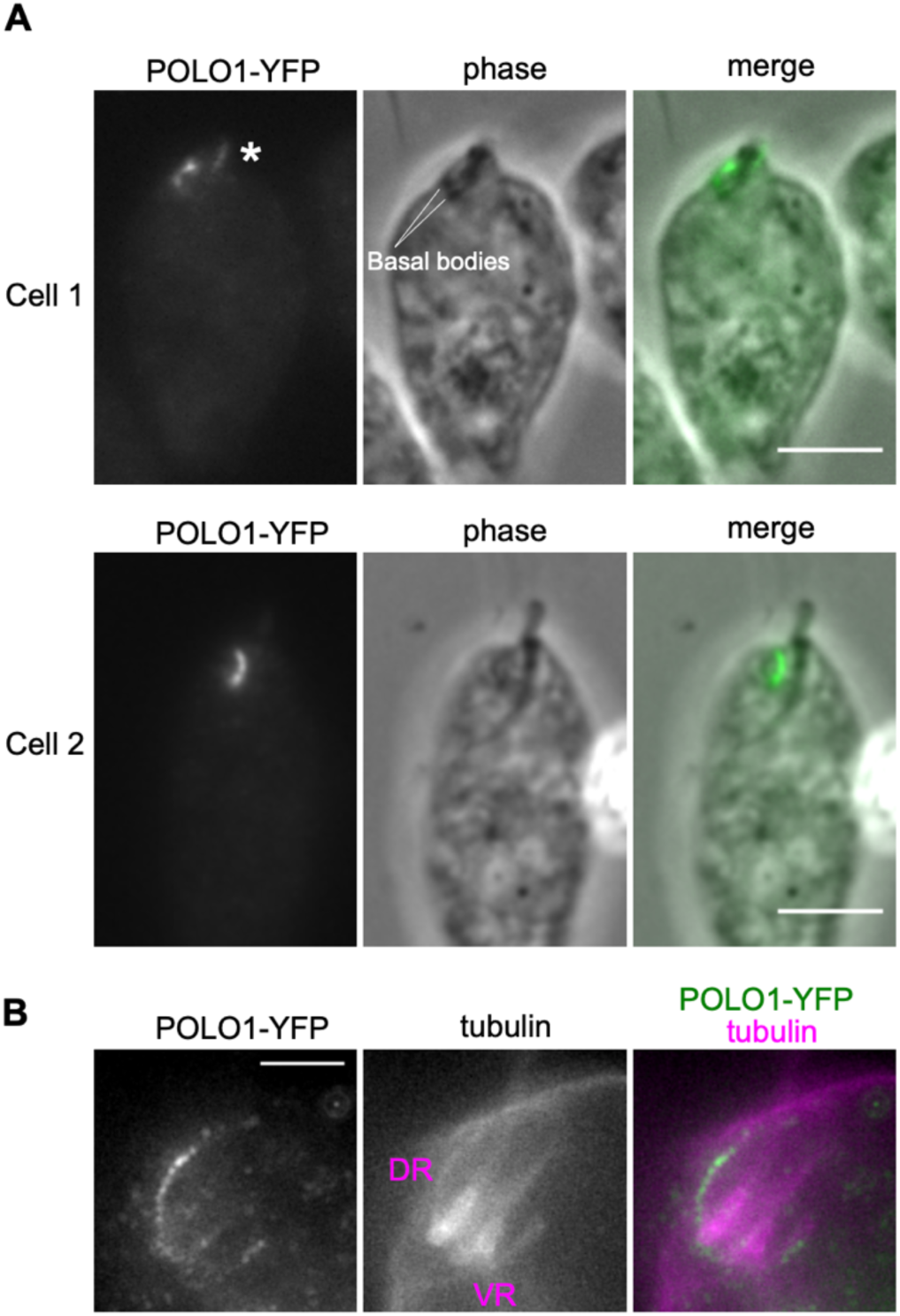
POLO1 localizes at the dorsal root. (A) POLO1-YFP signal is found near one of the two basal bodies and at an unknown location (asterisk). (B) U-ExM shows that POLO1 localizes at the DR. Scale bars, 5 µm. Cell line, DP10: POLO1-YFP.

**Figure 4.**
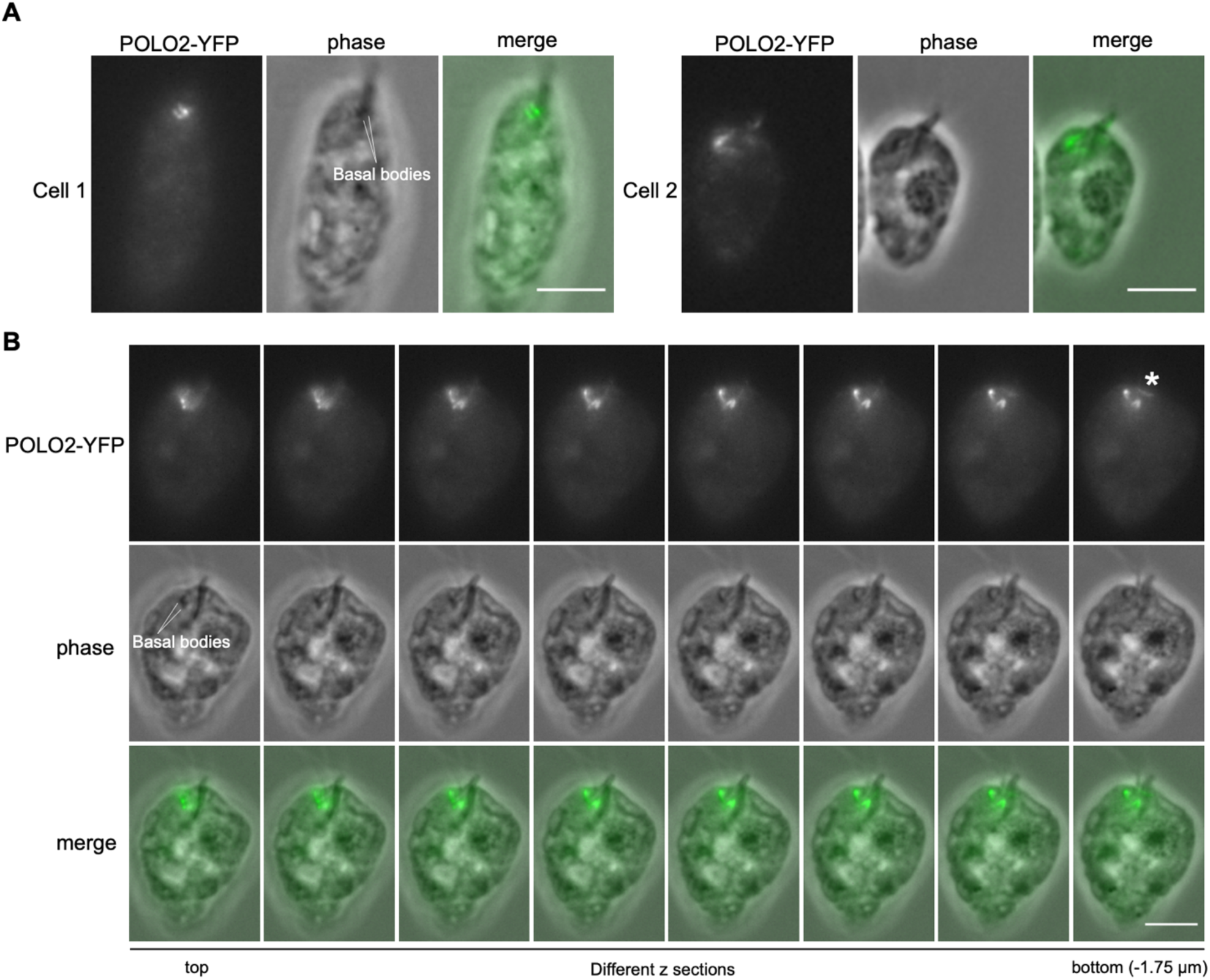
POLO2 localizes at and near basal bodies. (A) Two examples of cells expressing POLO2-YFP showing localization near both basal bodies, which may correspond to the DR and IR. (B) Z sections show a complex localization pattern of POLO2 at and near basal bodies, as well as at an unknown location (asterisk). Scale bars, 5 µm. Cell line, DP11: POLO2-YFP.

### KMP11A and PFR2

Kinetoplastid Membrane Protein-11 (KMP11) is a ∼10kDa protein that is highly conserved among kinetoplastids (Tolson et al., 1994). In *T. brucei*, it localizes at basal bodies and the flagellum (Li and Wang, 2008). Although it was initially thought that KMP11 is exclusive to kinetoplastids, its homologs are present in diplonemids (Valach et al., 2023). Four KMP11 genes are present in the *D. papillatum* genome, termed KMP11A (DIPPA_12089), KMP11B (DIPPA_12088), KMP11C (DIPPA_12130), KMP11D (DIPPA_12062) herein. KMP11A has 87% sequence identities to the *T. brucei* KMP11 protein, showing an extremely high level of similarity. KMP11A-YFP signal was found at basal bodies, part of flagella, apical papilla, and MTR, as well as widely at the feeding apparatus including the peripheral microtubule band (PMB) (Figure 5A, B). It is noteworthy that a ring-like structure was observed in one region of the cytopharynx (Figure 5B).

**Figure 5.**
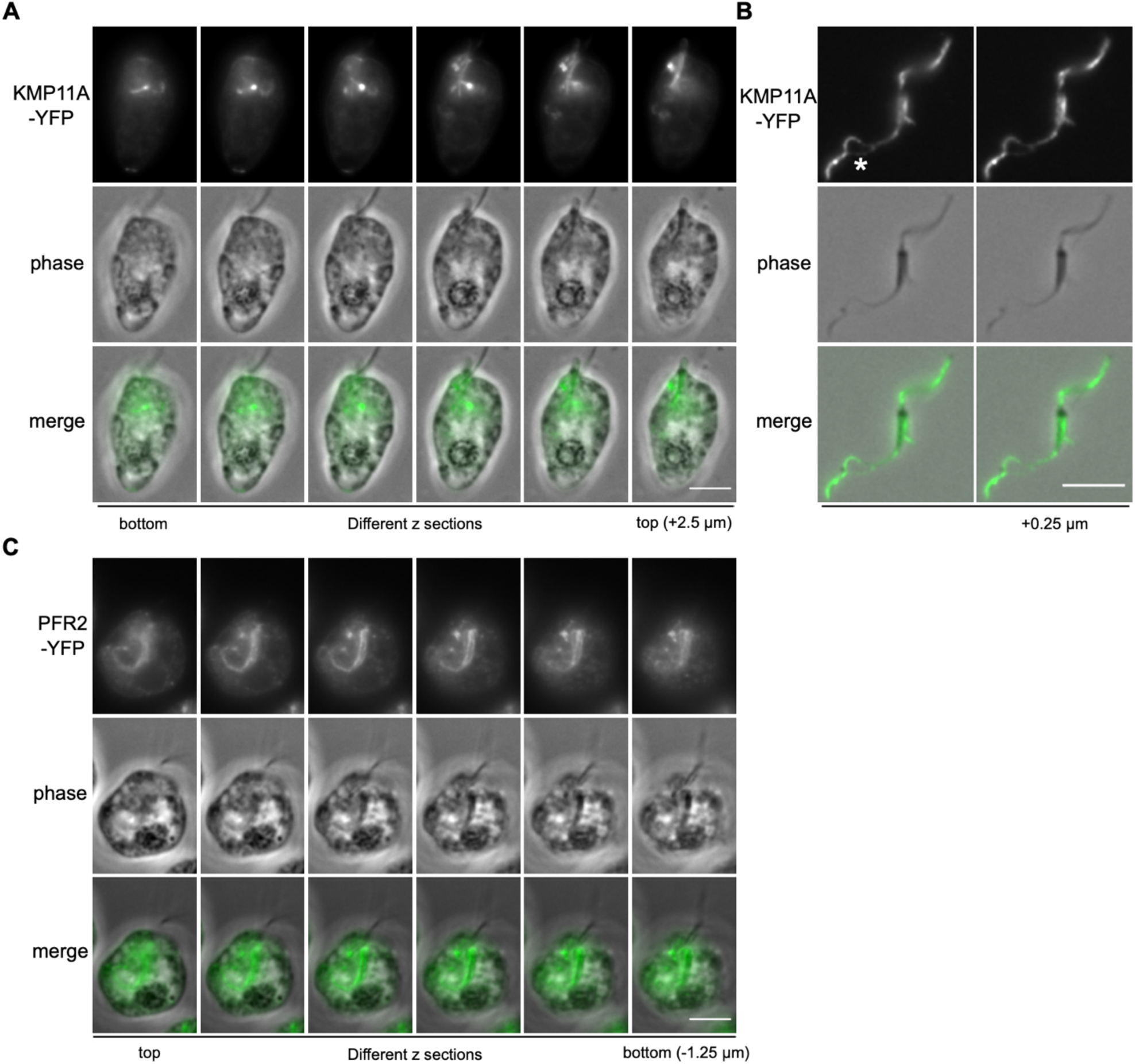
KMP11A and PFR2 localize at basal bodies and around the feeding apparatus. (A, B) Cells expressing KMP11A-YFP without (A) or with (B) detergent treatment were imaged. A loop structure is visible in the cytopharynx of detergent treated cells (asterisk). (C) Cells expressing PFR2-YFP shows a similar, but less defined, signal as KMP11. Additional images are shown in Figure S1A and S1B. Scale bars, 5 µm. Cell lines, DP127: KMP11A-YFP, DP79: PFR2-YFP.

The paraflagellar rod is a paracrystalline filamentous network that runs alongside the axoneme. In *T. brucei,* Paraflagellar Rod Protein 1/2 (PFR1/2) are major components of the paraflagellar rod (Portman and Gull, 2010). Interestingly, although *D. papillatum* lacks a paraflagellar rod (Porter, 1973), their homologs are present in the genome. We found that *D. papillatum* PFR2 (DIPPA_04871, which shares 77% and 69% identities to *T. brucei* PFR2 and PFR1, respectively) has a similar localization pattern as KMP11A, although the signal looked less defined (Figure 5C). No signal was found in detergent extracted cells.

### BILBO1 homologs localize at cytoskeletal structures

BILBO1 is a component of the flagellar pocket collar in *T. brucei* (Bonhivers et al., 2008; Zelená et al., 2025). *T. brucei* has at least four proteins that have similarities to BILBO1, all of which contain a ubiquitin-like N-terminal domain (NTD) (Vidilaseris et al., 2020). Although BILBO1-like proteins were thought to be kinetoplastid-specific, a large number of homologs are present in diplonemids. *D. papillatum* has 76 proteins that have similarities to the BILBO1 NTD as well as at least 10 proteins that have similarities in other regions of BILBO1 (see Methods). We suggest to call them BILBOD1–86 (File S1). BILBOD10-YFP had a complex localization pattern, including at the PMB, a part of cytopharynx, and the connection between cytopharynx and MTR (Figure 6A-C). In contrast, BILBOD26-YFP showed circle signals of unknown locations as well as signals around the feeding apparatus (Figure 6D). It will be interesting to examine the localization of other BILBOD proteins.

**Figure 6.**
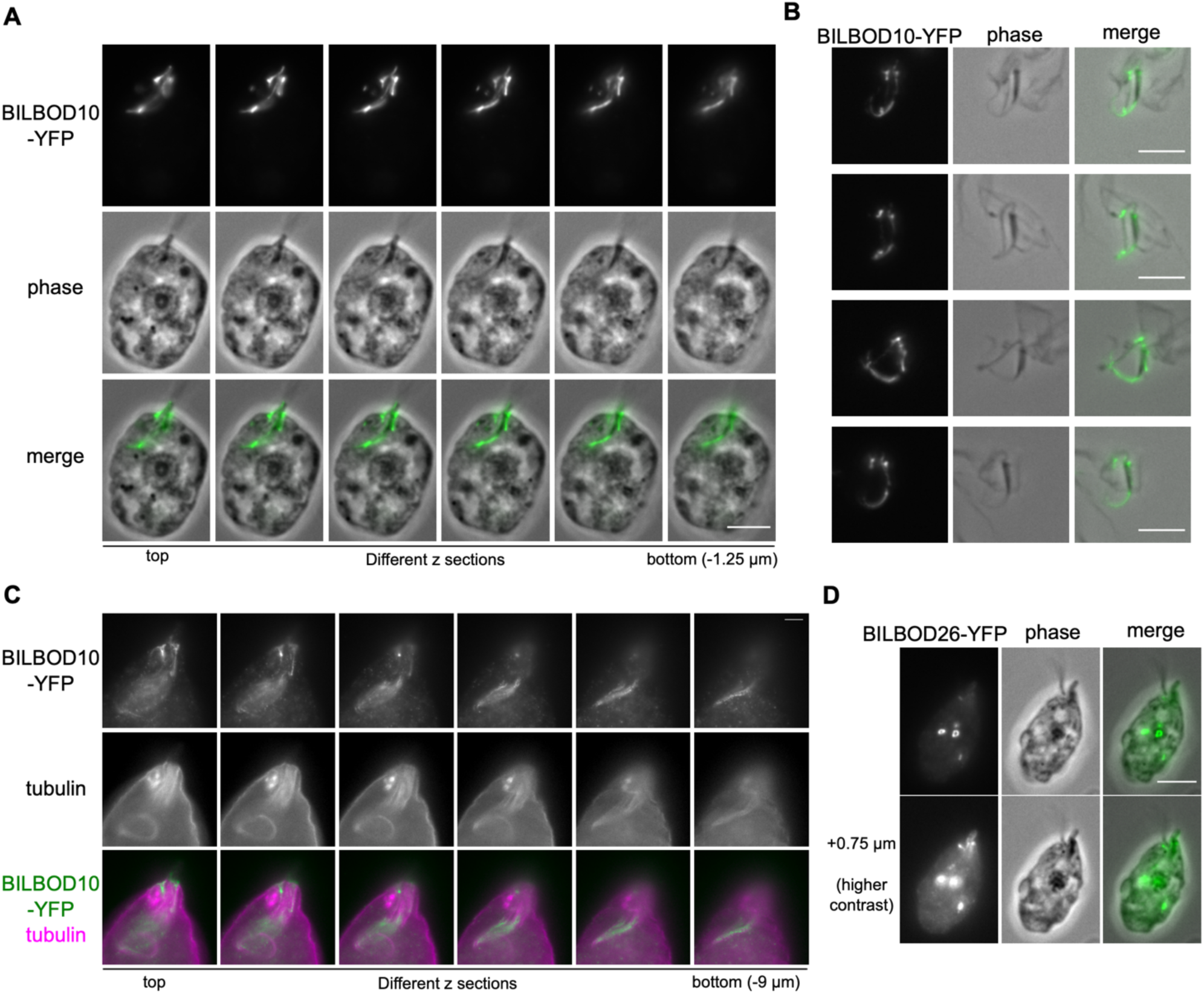
Two BILBO1 homologs localize near the feeding apparatus in *D. papillatum.* (A, B) BILBOD10 has a complex localization around the feeding apparatus. Cells expressing BILBOD10-YFP without (A) or with (B) detergent treatment were imaged. (C) U-ExM confirms the localization of BILBOD10 at the feeding apparatus area. (D) BILBOD26-YFP shows circle signals in the cytoplasm as well as a weaker signal around the feeding apparatus. Additional images for BILBOD10 are shown in Figure S1C, S1D. Scale bars, 5 µm. Cell lines, DP100: BILBOD10-YFP, DP99: BILBOD26-YFP.

### PTP2 localizes at apical papilla and PML

Protein tyrosine phosphatases (PTPs) are a diverse family of enzymes that remove phosphate groups from tyrosine residues on proteins, acting as crucial regulators of cell signaling, proliferation, and differentiation (Tautz et al., 2013). There are two PTPs in *D. papillatum*, termed PTP1 (DIPPA_30009) and PTP2 (DIPPA_18426) herein. We found that PTP2 localizes at the apical papilla, MTR, and PML (Figure 7A-C). These images reveal that the PML contacts the side of MTR as well as the tip of MTR.

**Figure 7.**
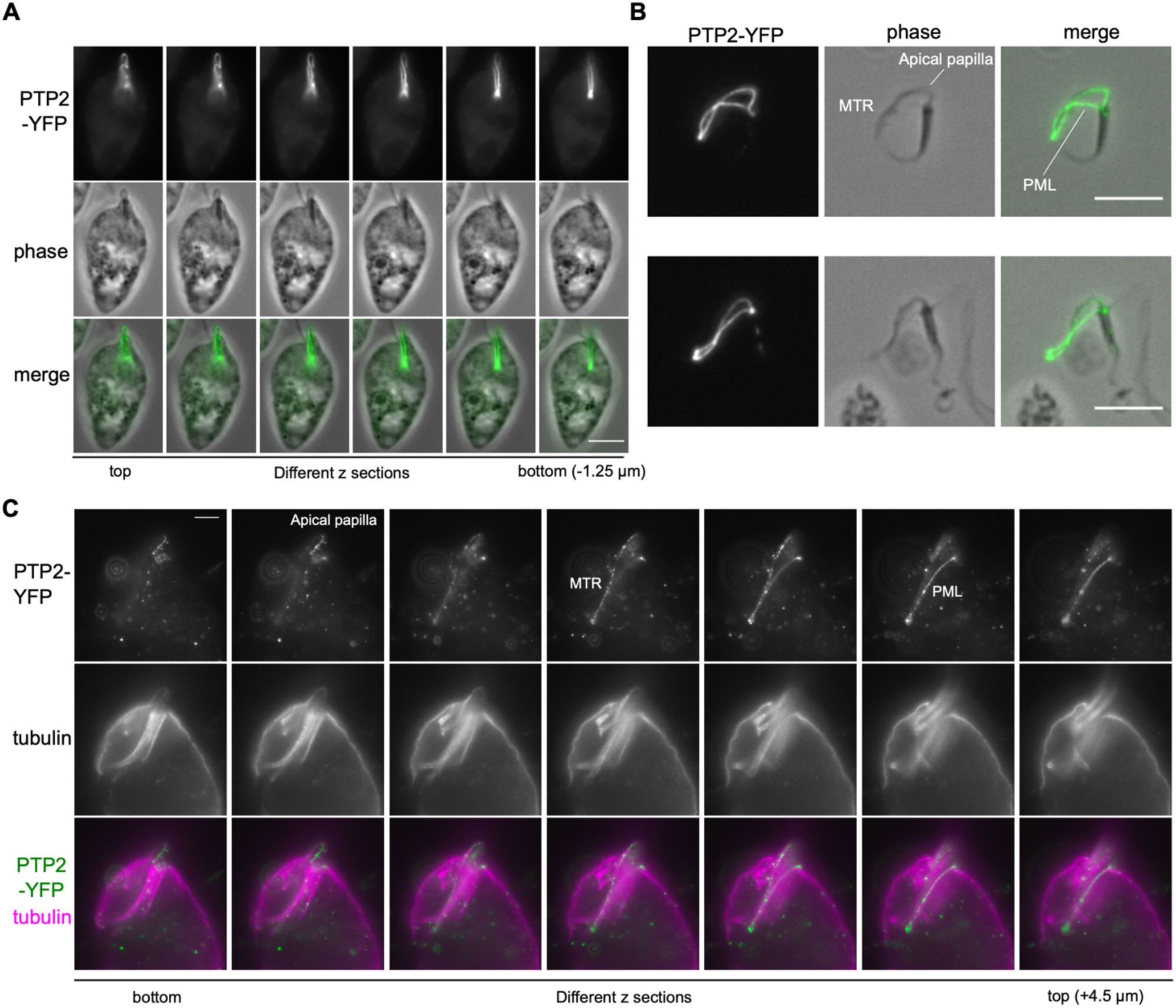
PTP2 localizes at apical papilla and PML. (A, B) PTP2 localizes at apical papilla (including MTR) and PML. Cells expressing PTP2-YFP without (A) or with (B) detergent treatment were imaged. (C) U-ExM of PTP2-YFP. Additional images are shown in Figure S2. Scale bars, 5 µm. Cell line, DP98: PTP2-YFP

### Other components that localize at feeding apparatus

We identified 8 additional components that localize at or near the feeding apparatus. KIFC1 (DIPPA_32533), a C-terminal kinesin motor protein, localizes at the MTR as well as basal bodies (Figure 8A), while a putative CENP-E protein (Benz et al., 2024) localizes near the tip of the MTR or the PML (Figure 8B). DIPPA_09231 is a highly degenerate kinesin-like protein that localizes at the bottom of the cytopharynx (Figure 8C), named KinesinWW herein because it has a WW domain, a protein-protein interaction domain that binds proline-rich motifs (Kay et al., 2000). KinesinWW lacks key residues required for a functional kinetochore motor (e.g. P-loop, Switch I, Switch II), suggesting that it is not a functional motor. Although it is highly conserved in diplonemids, putative orthologs of KinesinWW that have a WW domain are found only in prokinetoplastids (File S2).

**Figure 8.**
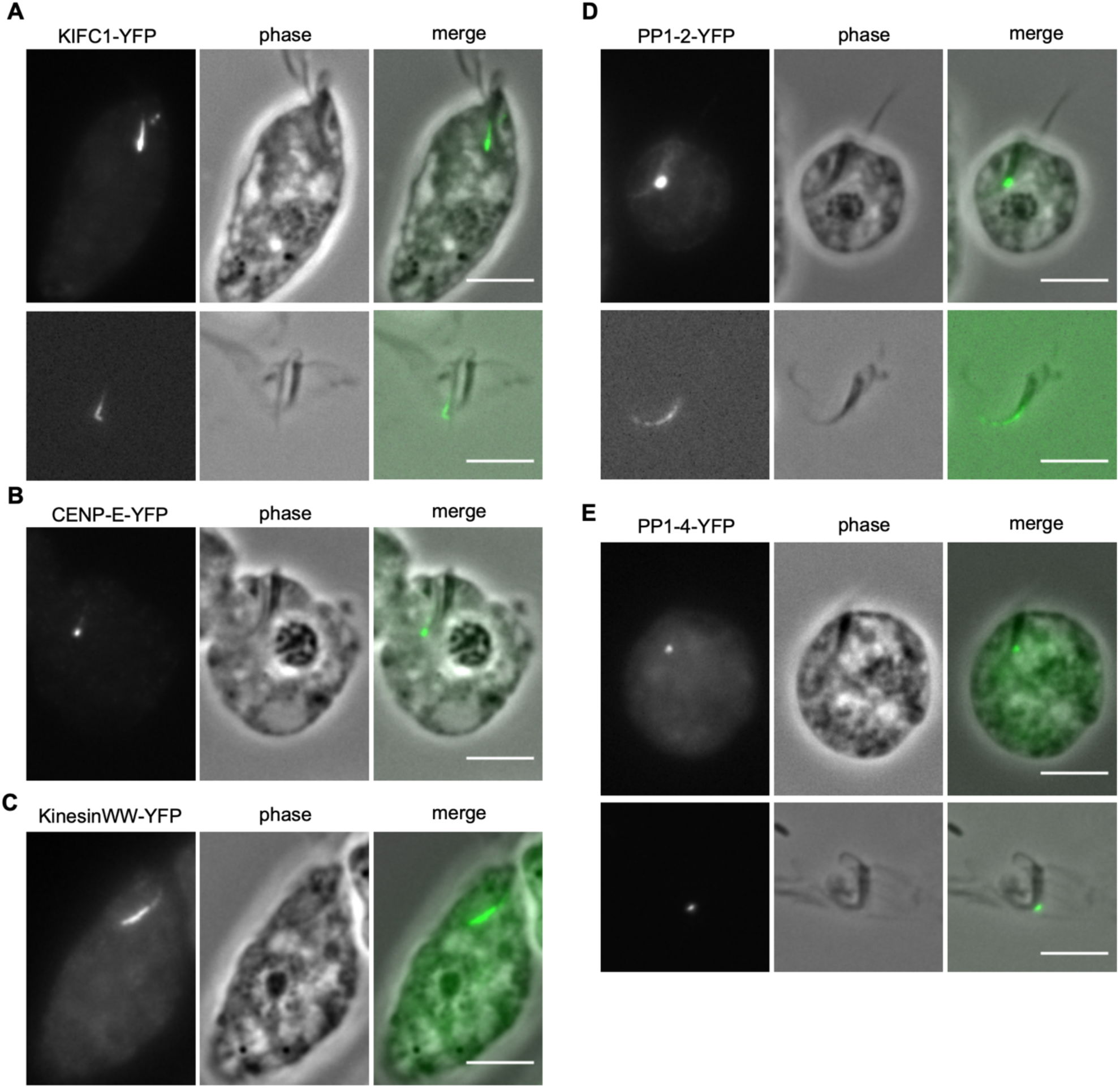
Identification of six proteins that localize to the feeding apparatus. (A) KIFC1-YFP localizes at basal bodies and the MTR. Top: non-treated cell, bottom: detergent-treated cell. (B) CENP-E-YFP localizes near the tip of the MTR. (C) KinesinWW-YFP localizes at the bottom of the cytopharynx. (D) PP1-2-YFP localizes at the region of cytopharynx where PMB terminates as well as cytopharynx. Top: non-treated cell, bottom: detergent-treated cell. (E) PP1-4-YFP shows signal at the region of cytopharynx where PMB terminates. Top: non-treated cell, bottom: detergent-treated cell. Scale bars, 5 µm. Cell lines, DP97: KIFC1-YFP, DP31: CENP-E-YFP, DP82: KinesinWW-YFP DP24: PP1-2-YFP, DP123: PP1-4-YFP.

*D. papillatum* has five homologs of protein phosphatase 1, termed PP1-1 (DIPPA_07735), PP1-2 (DIPPA_07733), PP1-3 (DIPPA_25219), PP1-4 (DIPPA_29493), and PP1-5 (DIPPA_21477) herein. PP1-2 localizes at the region of cytopharynx approximately where the PMB terminates as well as the bottom of the cytopharynx (Figure 8D). By contrast, PP1-4 localizes only at the region of cytopharynx where the PMB terminates (Figure 8E).

Finally, by tagging proteins that are conserved among diplonemids but have no annotation, we identified three components of the feeding apparatus. The protein encoded by DIPPA_13090 localizes at the MTR region (Figure 9A). We therefore named it MTR1. Kinetoplastids have MTR1 homologs (File S3), and the *T. brucei* homolog (Tb927.10.630) localizes at the hook complex (Billington et al., 2023). The protein encoded by DIPPA_22779 localizes at the PML (Figure 9B, C), so we named it PML1. Besides diplonemids, its putative homologs are found only in prokinetoplastids (File S4). PML1 has a caspase-like domain in the N-terminal part of the protein. The protein encoded by DIPPA_20809 specifically localizes at the apical papilla (Figure 9D, E). We named the protein APL1. Its homologs are present in free-living kinetoplastids and trypanosomatids that have a cytostome-cytopharynx complex (*T. cruzi* and *Crithidia*) (File S5), but not in *T. brucei* or *Leishmania*.

**Figure 9.**
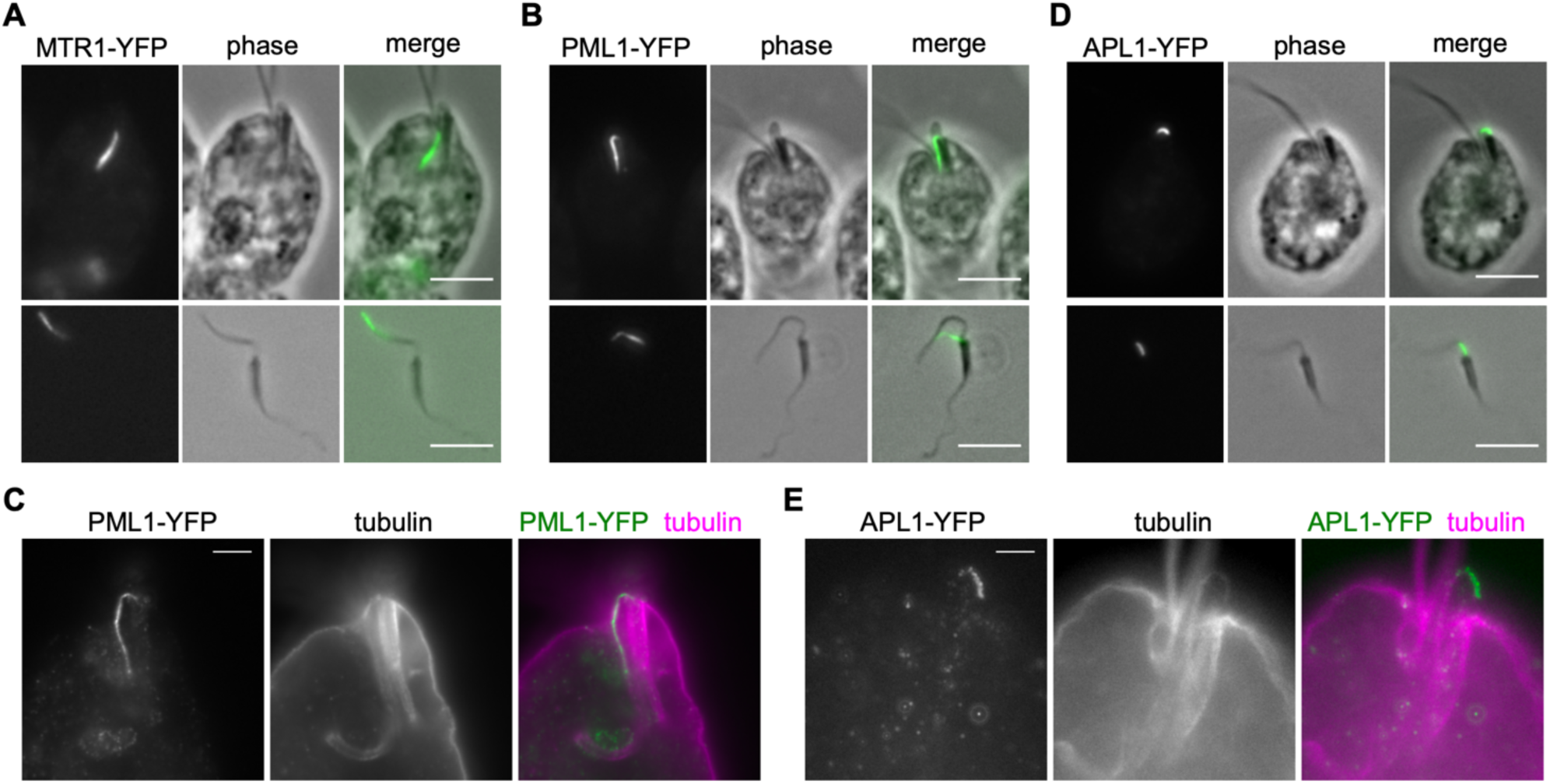
Proteins that localize at PML or apical papilla. (A) MTR1-YFP localizes at the MTR. Top: non-treated cell, bottom: detergent-treated cell. (B) PML1-YFP has signal at the PML. Top: non-treated cell, bottom: detergent-treated cell. (C) U-ExM of PML1-YFP. (D) APL1-YFP specifically localizes at the beginning of the apical papilla. (E) U-ExM of APL1-YFP. Scale bars, 5 µm. Cell lines, DP69: MTR1-YFP, DP119: PML1-YFP, DP61 APL1-YFP.

## Conclusions

Determining the composition of macromolecular structures is a key step towards their mechanistic understanding. So far very little is known about the components of the cytostome-cytopharynx complex found in flagellates and ciliates. Experimental tractability of *D. papillatum* makes it an attractive model to identify molecular components of the feeding apparatus in diplonemids, a highly abundant flagellates in the world’s ocean. Furthermore, some components are conserved in disease-causing kinetoplastid parasites such as *T. cruzi* whose cytostome-cytopharynx complex could be a drug target. Immunoprecipitation of YFP-tagged proteins reported in this work, followed by mass spectrometry has a potential to identify many additional components of the feeding apparatus in *D. papillatum* and beyond.

## Materials and Methods

### Cells, plasmids and primers

Plasmids and primer sequences used in this study are provided in Table S1. C-terminal YFP-tagging constructs were made as previously described using pBA3294 (Akiyoshi et al., 2025). Briefly, two ∼2 kb homology arms amplified from genomic DNA by PCR using KOD one polymerase (Merck) were inserted into pBA3294 cut with *Pac*I and *Asc*I with a unique site in between the two PCR fragments (typically *Not*I; if *Not*I was not unique, *Sbf*I or *Spe*I were used as indicated). Plasmids were validated by nanopore whole plasmid sequencing (Plasmidsaurus). As previously observed in our attempts to tag other genes (Akiyoshi et al., 2025), there were occasional mismatches between nanopore sequencing results and expected plasmid sequences, especially in repetitive sequences in 3′UTR regions.

### Diplonema cultures and YFP-tagging of genes at the endogenous locus

All cell lines used in this study were derived from *Diplonema papillatum* (ATCC 50162) and are listed in Table S1. Cells were grown at 27 °C in artificial sea water containing 36 g/L Instant Ocean Sea Salt (Instant Ocean), 1 g/L trypton (Formedium, TRP01), and 1% fetal bovine serum (Merck, F9665) in vented flasks. For transfection of C-terminal YFP-tagging constructs, approximately 5 to 10 µg plasmids were linearized by *Not*I or other enzymes, cleaned up by ethanol precipitation, resuspended in transfection reagent (Ingenio Electroporation Kit for the EZporator Electroporation System, Cambridge Biosciences), and transfected into 2–3 x 10^7^ cells using Amaxa Nucleofector IIb (Lonza Bioscience) as previously described (Faktorová et al., 2026). Transfected cells were selected by addition of 75 µg/mL G418 (Merck).

### Microscopy

To observe native YFP signals without detergent extraction, cells fixed in formaldehyde were imaged as previously described with minor modifications (Akiyoshi et al., 2025). Briefly, 1–2 mL cell culture was centrifuged at 2000 g for 3 min. Cells were fixed by 4% formaldehyde solution (Life technologies, 28906) diluted in PBS for 5 min, rinsed with PBS twice, resuspended in a small volume (∼10 µL) of DABCO mounting media (1% w/v 1,4-diazabicyclo[2.2.2]octane, 90% glycerol, 50 mM sodium phosphate pH 8.0) with 100 ng/mL DAPI, and mounted onto glass slides. To observe YFP signals on detergent-treated cells, 2 mL cell culture was centrifuged at 2000 g for 3 min. The pellet was treated with PEME (100 mM PIPES-NaOH 6.9, 2 mM EGTA, 1 mM MgSO_4_ and 0.1 mM EDTA) with 1% NP-40 and immediately centrifuged at 2000 g for 3 min, followed by fixation and slide preparation as described above. Images were captured on an Axioimager.Z2 microscope (Zeiss) installed with ZEN using a Hamamatsu ORCA-Flash4.0 camera with 63x objective lenses (1.40 NA). Typically, 15–25 z sections covering 3–6 µm were collected. Images were analyzed in ImageJ/Fiji (Schneider et al., 2012). Figures were made in Inkscape (version 1.4, https://inkscape.org/).

### U-ExM

Ultrastructure expansion microscopy of *D. papillatum* was carried out by adapting protocols used in *T. brucei* (Amodeo et al., 2021; Gorilak et al., 2021). 1 mL cell culture was centrifuged at 2000 g for 3 min. Cells were fixed in 4% formaldehyde solution diluted in PBS for 5 min, followed by centrifugation. The pellet was resuspended in 500 µL of fixative solution (0.7% formaldehyde, 1% acrylamide (Merck, A4058) in PBS). Cells were settled onto a 12 mm coverslip (Thermo, No. 1.5, CB00120RA120MNZ0) overnight at room temperature. Gelation was performed on a glass slide wrapped with parafilm on a flat metal surface in ice bucket. 50 µL of the monomer solution (19% sodium acrylate (Fluorochem, 461652), 10% acrylamide and 0.1% N, N’-methylene-bis-acrylamide (Bis) (Merck, M7256) in PBS) chilled on ice was quickly mixed with 0.25 µL of TEMED (MP Biomedical, 04805613) and 2.5 µL of 10% ammonium persulfate (Thermo, 17874), and put on the parafilm. Immediately after this, the coverslip which cleared excess liquid (with the cells facing down) was placed on the monomer solution and incubated for 5 min. Then the slide was moved to a wet chamber and incubated at 37°C for 30 min to promote gel polymerization.

The parafilm was cut into pieces to move the gel/coverslip to a 6-well plate containing 2 mL of denaturation buffer (50 mM Tris pH9.0, 200 mM SDS, 200 mM NaCl) to promote detachment of the gel from the coverslip. The gel was then transfer to a 1.5 mL tube containing 1 mL of denaturation buffer, and incubated at 95°C for 60 min. Then the gel was moved into a 12 cm square petri dish containing 10–20 mL MilliQ water and expanded by 3x changes into fresh MilliQ water. After checking the expansion factor (typically 4.0–4.2), the gel was cut into smaller pieces (∼1 cm x 1 cm square) for immunostaining. The gel piece was moved to a 6-well plate and shrunk in PBS for 30 min. Gels were incubated with 600 µL of primary antibodies (rabbit polyclonal anti-GFP antibody (OriGene, TP401, 1 mg/mL, used at 1:500 dilution) and mouse monoclonal anti-alpha tubulin antibody (Merck, T5168, used at 1:200 dilution) diluted in PBS with 1% BSA) overnight on a rocker at room temperature. Gels were washed with 4 mL PBS for 15 min, three times, followed by incubation with secondary antibodies (goat Alexa Fluor 488 anti-rabbit IgG (H+L) (Thermo, A11008, used at 1:200 dilution) and goat Alexa Fluor 568 anti-rabbit IgG (H+L) (Thermo, A11004, used at 1:200 dilution) diluted in PBS with 1% BSA) overnight in dark on a rocker at room temperature. Gels were washed with PBS for 10 min, three times. Gels were expanded with 4 mL MilliQ with 0.2% propyl gallate (Merck, P3130) for 15 min, three times. Gels were mounted onto glass-bottom chambers 35 mm (No. 1.5, Thistle Scientific, IB-81218-200) coated with poly-L-lysine (Merck, P4707). Images were captured on an Axioimager.Z2 microscope (Zeiss) installed with ZEN using a Hamamatsu Flash 4 sCMOS camera with 63x objective lenses (1.40 NA). Typically, 167 z sections covering 25 µm were collected.

### Positive Staining and TEM

Carbon Films on 400 mesh gold grids (Agar Scientific, AGS160A4) were glow-discharged for 2 min (PELCO easiGlow, 25 mA current) before being deposited carbon-side down on 20 μL 0.01% Poly-L-Lysine (Sigma, P4707) droplets and incubated for 20 min at room temperature. Grids were then washed 3x by dipping in 30 μl droplets of MilliQ water, residual water removed (in this and subsequent steps by dabbing the side of the grids with Whatman Grade 1 filter paper (Cytiva)), and placed carbon-side up on a glass slide. *Diplonema papillatum* cell culture was then added to the slide and incubated at room temperature for 75 min. Cell culture was then removed, and grids were washed 3x in 30 μL droplets of PEME (100 mM PIPES-NaOH 6.9, 2 mM EGTA, 1 mM MgSO_4_ and 0.1 mM EDTA). Residual PEME was removed before adding 6 μL 1% NP40 (diluted with PEME from 10% NP40 stock solution in MilIiQ water) and incubating for 1 min at room temperature. Then, residual liquid was removed before grids were deposited on 30 μL droplets of 2.5% glutaraldehyde (25% stock solution diluted with PEME) and incubated for 20 min at room temperature. Grids were then washed 5x in 30 μL PEME droplets and residual PEME removed, before 6 μL 2% uranyl acetate was added to the grids and incubated for 30 s. Then, they were washed 5x in 30 μL MilliQ water droplets, residual water removed, and grids deposited onto filter paper in a glass petri dish and left to dry overnight at room temperature. TEM images were acquired using a 120kV JEOL TEM-1400 Plus with a Gatan OneView camera using Digital Micrograph version 3.61.4719.0. The images in Figure 1 were both taken with an exposure time of 1 s. The image in Figure 1A was taken at a magnification of 5kX with -4.813 μm defocus and the image in Figure 1B was taken at a magnification of 4kX with -9.626 μm defocus.

### Bioinformatics analysis

Protein sequences were retrieved or accessed from UniProt (UniProt Consortium, 2019), TriTryp (Aslett et al., 2010), EukProt (Richter et al., 2022) or published studies (Valach et al., 2017; Butenko et al., 2020; Kaur et al., 2020; George et al., 2022; Valach et al., 2023). Homologs were aligned using with MAFFT (L-INS-i, version 7) (Katoh et al., 2019) and visualized with the Clustalx coloring scheme in Jalview (version 2.11) (Waterhouse et al., 2009). Homology searches were done using BLAST in TriTryp (Aslett et al., 2010), EukProt v3 BLAST server (Richter et al., 2022), or hmmsearch using manually prepared HMM profiles (HMMER version 3.0) (Eddy, 1998). Domains were searched using HHpred (Zimmermann et al., 2018) or Foldseek (van Kempen et al., 2024) of AlphaFold2 structural models (Jumper et al., 2021; Abramson et al., 2024; Benz et al., 2024).

BILBO1 homologs in *D. papillatum* were identified as follows. We first searched BILBO1 homologs in diverse kinetoplastids (*Bodo saltans*, *Trypanosplasma borreli*, *Azumiobodo hoyamushi*, *Papus ankaliazontas*, *Apiculatamorpha spiralis*). We then performed multiple sequence alignment together with *T. brucei* BILBO1 using MAFFT (Katoh et al., 2019), from which a BILBO1-NTD hidden Markov model was created (corresponding to 1–123 residues of TbBILBO1) using HMMER (Eddy, 1998). We then performed hmmsearch against *D. papillatum* protein database (Valach et al., 2023; Gray et al., 2025), identifying 76 hits (BILBOD1–76). Ten additional homologs (BILBOD77–86) were identified using full-length BILBO1 protein sequences.

## Acknowledgments

This work was supported by funding for the Wellcome Discovery Research Platform for Hidden Cell Biology (226791) and we gratefully acknowledge support from the Microscopy/Structural Biology cores. Bungo Akiyoshi was supported by a Wellcome Discovery Award (227243/Z/23/Z).

## Author contributions

G.C. made transgenic lines and performed imaging. S.J.P.T. and F.C.-S. performed TEM. M.I. optimized expansion microscopy for *D. papillatum*. B.A. made transgenic lines, performed imaging and expansion microscopy, and wrote the manuscript.

## Rights retention

This research was funded in whole, or in part, by the Wellcome Trust (227243/Z/23/Z). For the purpose of open access, the author has applied a CC BY public copyright licence to any Author Accepted Manuscript version arising from this submission.

## Acronyms

IR: intermediate root
VR: ventral root
DR: dorsal root
MTR: reinforcing microtubules
PML: parallel microtubule loop
PMB: peripheral microtubule band

## Supplementary Information

**Supplemental_files.zip contains the following files:**

**Table S1: List of D. papillatum cell lines, plasmids, and primer sequences used in this study**

**File S1: Protein sequences of BILBOD1–86**

**File S2: Multiple sequence alignment of KinesinWW**

**File S3: Multiple sequence alignment of MTR1**

**File S4: Multiple sequence alignment of PML1**

**File S5: Multiple sequence alignment of APL1**

## Supplementary Figures

**Figure S1.**
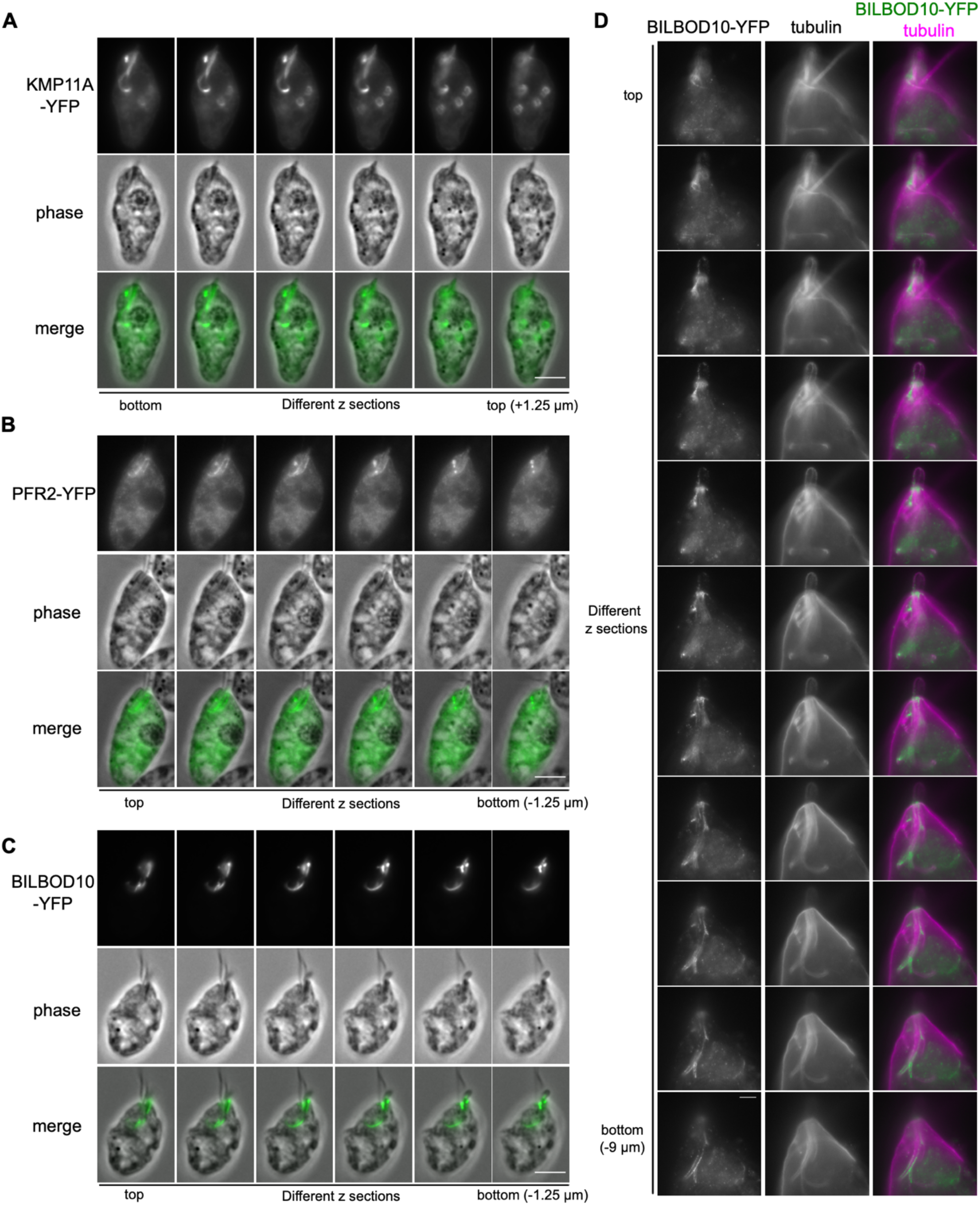
Additional images of KMP11A, PFR2, and BILBOD10. (A) KMP11A-YFP. (B) PFR2-YFP. (C) BILBOD10-YFP. (D) U-ExM of BILBOD10-YFP. Scale bars, 5 µm.

**Figure S2.**
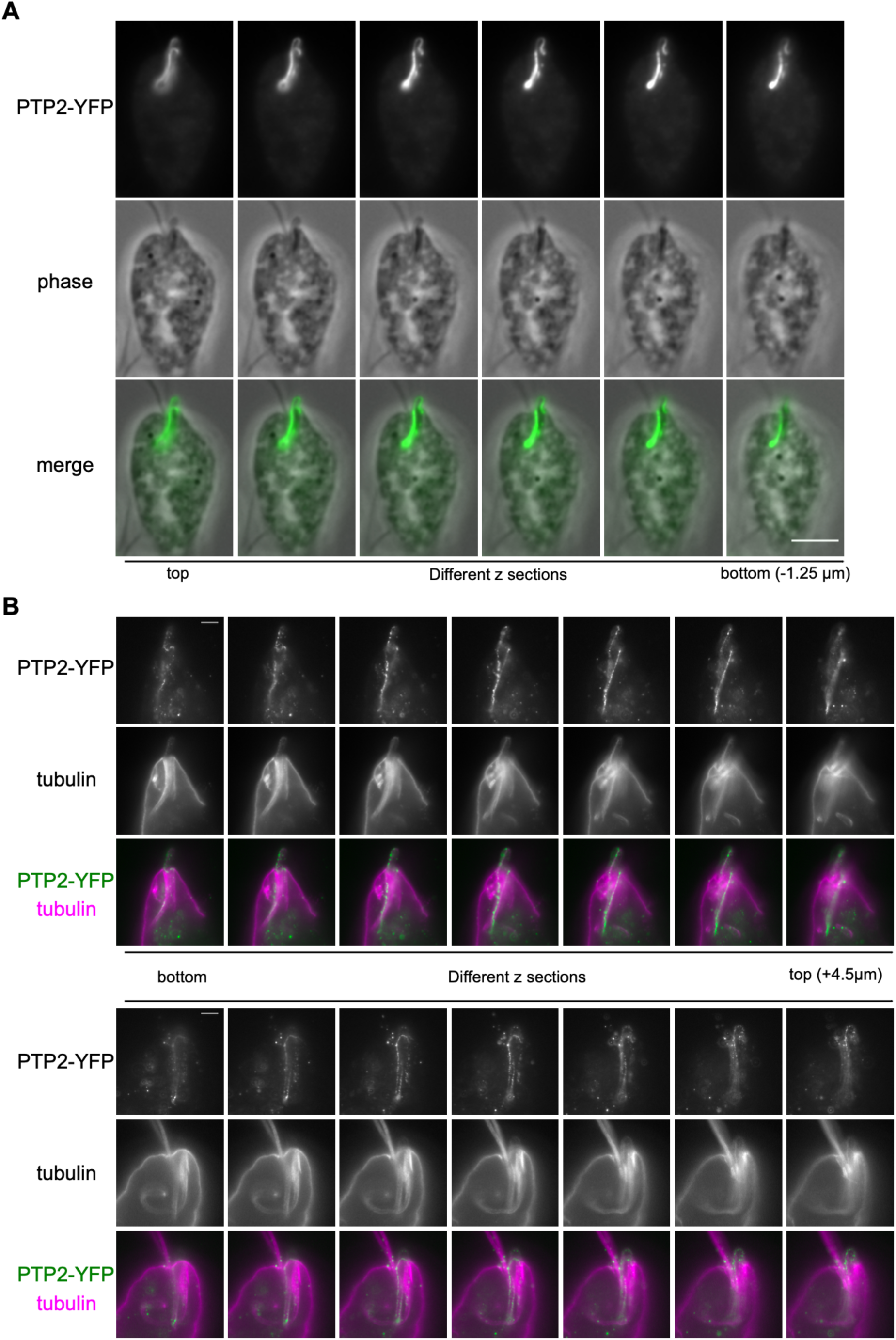
Additional images of PTP2-YFP. (A) A cell expressing PTP2-YFP. (B) U-ExM of PTP2-YFP. Scale bars, 5 µm.

